# GBRAP: A Comprehensive Database and Tool for Exploring Genomic Diversity Across All Domains of Life

**DOI:** 10.1101/2024.10.28.620620

**Authors:** Sachithra Kalhari Yaddehige, Chiara Vischioni, Leonardo Alberghini, Michele Berselli, Cristian Taccioli

## Abstract

Evolutionary studies require extensive examination of genomic information across all domains of life. Despite the availability of a large number of genomes through GenBank, the effective visualisation or comparison of the information they contain is challenging due to many reasons including their size. We introduce GBRAP, a comprehensive software tool to analyse GenBank files, and an online database housing an extensive collection of carefully curated, high-quality genome statistics for all the organisms available in the RefSeq database of NCBI. Users can either directly search, or select from pre-categorized groups, the organisms of their choice and retrieve data, which eventually output as tables containing more than 200 columns of useful genomic information (Base counts, GC content, Shannon Entropy, Codon Usage etc.) separately calculated for different genomic elements (eg. CDS, Introns, tRNA, rRNA, ncRNA, etc.). The data are independently displayed (if applicable) for each chromosomal, mitochondrial, plastid, or plasmid sequence. All the data can be visualised on the database or downloaded as CSV or Excel files. The GBRAP database is free to access without any registration and is publicly available at <GBRAP database link will be available here>.

## Introduction

Understanding the evolutionary relationships across different life kingdoms requires extensive examination and comparison of genomic data. The vast array of genomes available through the National Center for Biotechnology Information (NCBI) offers a unique opportunity to explore these relationships. However, the mere availability of genomes does not ensure the effective interpretation or comparison of the information they contain, making it necessary to transform these genetic data into numerical forms to facilitate better understanding.

Base frequencies, GC content, or genome size are commonly used to study evolutionary relationships among organisms (Bohlin and H-O Pettersson 2019). Base frequencies represent the relative abundance of each nucleotide (adenine, thymine, cytosine, and guanine), and provide information about the genomic composition while GC content which is the proportion of guanine (G) and cytosine (C) bases in a sequence, can provide information about the splice site usage and structural stability of RNA sequences (Zhang et al. 2011). Codon usage is another well-known metric, referring to the frequency with which different codons are used in coding sequences. This can reveal biases in the genetic code that are shaped by evolutionary pressures, such as selection, mutation and genetic drift (Hershberg and Petrov 2008). Variations in codon usage can also provide insights into gene expression levels and the adaptation of organisms to their environments (Egelkrout et al. 2012; Arella et al. 2021).

Entropy as a measure of the randomness or disorder of a system is normally calculated in many forms (Arfken et al. 1984). From a genomics perspective, it can be used to quantify the diversity of DNA sequences of varying lengths (Koslicki 2011) or to quantitively describe the complexity of a sequence based on its information content (Li et al. 2019). For example, according to Shannon’s entropy, a sequence with ATCGAT can be described as having more information content compared to a sequence with AAAAAA (Shannon 1948). Another way to calculate entropy in numerical format is with Topological entropy which is able to characterize the complexity of a sequence (Koslicki 2011). Another less common yet important metric for comparative genomics is Chargaff’s second parity rule (PR2). It states that in each single strand of dsDNA (double-stranded DNA), the proportion of adenine (A) is approximately equal to thymine (T), and the proportion of cytosine (C) is approximately equal to guanine (G) (Rudner et al. 1968). Fariselli et al in 2021, introduced a new method to calculate a sequence’s adherence to PR2 as a numerical value where a sequence having a score of 1 perfectly follows PR2. Their analysis also demonstrated that PR2 provides insights into codon bias selection in the human genome (Fariselli et al. 2021).

These various genomic metrics allow researchers to perform comparative analyses between genomes to uncover evolutionary relationships and underlying mechanisms driving genomic diversity. Although calculating these metrics is straightforward for small genomes, interpreting them for large genomes remains challenging and time-consuming. This necessitates the development of advanced tools and databases capable of handling complex genomic data of any size and interpreting their characteristics accurately and efficiently. Nowadays, similar databases usually focus on a single group of organisms or a specific species (Shamimuzzaman et al. 2020; Roces et al. 2024), being somehow limited in their scope. Moreover, they typically include only a few data metrics, lacking detailed information at the level of individual chromosomes or single genomic elements (Subramanian et al. 2022).

To fill this gap, we developed GBRAP (Genome-Based Retrieval and Analysis Parser), a software tool proficient in parsing GenBank files. The data generated by GBRAP was then used to build an online database which houses and curates the information collected by the software. This repository, with easily accessible data of all the genomes available in RefSeq, yields valuable insights into genomic diversity, complexity, and base composition parity across all domains of life through one single platform.

### Novelty

GBRAP adds significant novelty to the panorama of genomics databases developed for comparative purposes. Our software tool can process and analyse a diverse range of genomic data, from small genomes like those of bacteria and viruses to larger ones belonging to amphibians, plants, and mammals. Additionally, the GBRAP database provides these data separately (where applicable) for each chromosomal, mitochondrial, plastid, or plasmid sequence across 49,330 organisms.

The types of genomic data available in GBRAP add further substantial value to the field, as they extend beyond basic information such as base counts, frequencies, and GC content to include some advanced genomic information such as topological entropy (Koslicki 2011), Shannon’s entropy (Shannon 1948), and Chargaff’s second parity rule (PR2) score (Fariselli et al. 2021) (see supplementary materials for further details about calculations). Altogether, these data will allow researchers to study the complex and evolutionary patterns between organisms from a new perspective. Indeed, by encompassing a broader range of genomic elements, GBRAP enables a more comprehensive and detailed understanding of genome organization and function, providing data not only for entire chromosomes but also for distinct genomic elements along with their strand-specific counts. To our knowledge, GBRAP is the first database and tool to offer such an extensive range of information through a single platform. Finally, it also extends the query target beyond coding sequences (CDS) and genes to include less-characterized elements like introns, tRNA, rRNA, and non-coding RNA (ncRNA). These elements offer valuable insights into regulatory mechanisms, structural roles, and evolutionary processes that are often overlooked when focusing solely on coding sequences. This inclusive approach ensures that researchers have access to critical information that can drive new discoveries in genomics and enhance our understanding of biological complexity and evolution.

Not just the quantity of data but GBRAP stands out in the quality of data as well. This high quality was achieved by applying several filtering mechanisms within the GBRAP script to extract data, which were developed after careful examination of the gbff files. First, the script is able to successfully filter out alternate loci and scaffolds, ensuring data extraction is exclusively from complete chromosomal and mitochondrial (and plastid, or plasmid) sequences. If the entire genome file is made of incomplete sequences, the organism is removed from the database. Furthermore, GBRAP can selectively identify one isoform per gene through several filters to prevent data redundancy. This level of intense filtering has not been addressed in other comparable databases.

### Implementation

GBRAP software tool was developed in Python 3 and is operated through the command line. The script only utilises standard Python libraries like math and regex which saves computational time and avoids the need for additional installations. As a result, GBRAP software can be efficiently executed even on a standard laptop without the need for specialized hardware or configurations. GenBank Flat Files (.gbff format) are used as inputs for the GBRAP tool. These files contain one or multiple sequences (chromosomal, mitochondrial/chloroplast, plastid, or plasmid) and their annotations. Using a Python script, we downloaded the gbff files of all the organisms with a reference or a representative genome available in the RefSeq repository of NCBI. A total of 64,111 gbff files (2.64 TB) belonging to 9 taxa were downloaded and used as inputs for our analyses.

The GBRAP script defines a class that features several functions to efficiently extract data from gbff files, filter the data, and calculate different genomic scores. These functions were carefully curated following thorough examinations of gbff files. For instance, we observed in gbff files, that some isoforms of CDS do not follow the general biological format of a coding sequence. Then we developed the ‘cds_selector’ function in the script to filter out such isoforms. It operates by first removing CDS isoform sequences that do not start with a start codon, and then eliminating isoforms that are not composed entirely of triplets of bases (not divisible by 3). If multiple isoforms persist, the function selects the longest among them to avoid data redundancy.

The time taken by the tool to analyse one genome varied depending on the size of the genome. The entire analysis was performed on a server with an Intel(R) processor, 36 processing cores, and 125 GB of RAM. Approximately, it took 2-4 seconds to analyse small genomes such as viruses or bacteria while it took about 20-30 minutes to analyse big genomes such as vertebrates (e.g Homo sapiens [T2T-CHM13]) or plants with some exceptions (∼1hr) for very large (>10Gb) genomes. The output was generated as comma-separated (CSV) files containing data across 219 columns (see supplementary material for details on all the columns) and rows equivalent to the number of chromosomes (plus mitochondrial/chloroplast, plastid, plasmid) the organism has. Once the analysis was completed for all the organisms, a SQL database containing data of 49,330 organisms was constructed using MySQL database management system.

### GBRAP Database

The GBRAP online database, developed using HTML and PHP, is a user-friendly and straightforward platform offering both visualizable and downloadable data, freely accessible without any registration. The homepage of GBRAP is displayed in Figure 1a, where users can either search for an organism by typing its scientific name in the search box and selecting it from the drop-down list or clicking on pre-categorized groups (Archaea, Bacteria, Mammals, Amphibians, etc.) to view a list of organisms within the chosen group. Either way, the user will be directed to the table of the chosen organism as displayed in Figure 1b).

**Figure 1:**
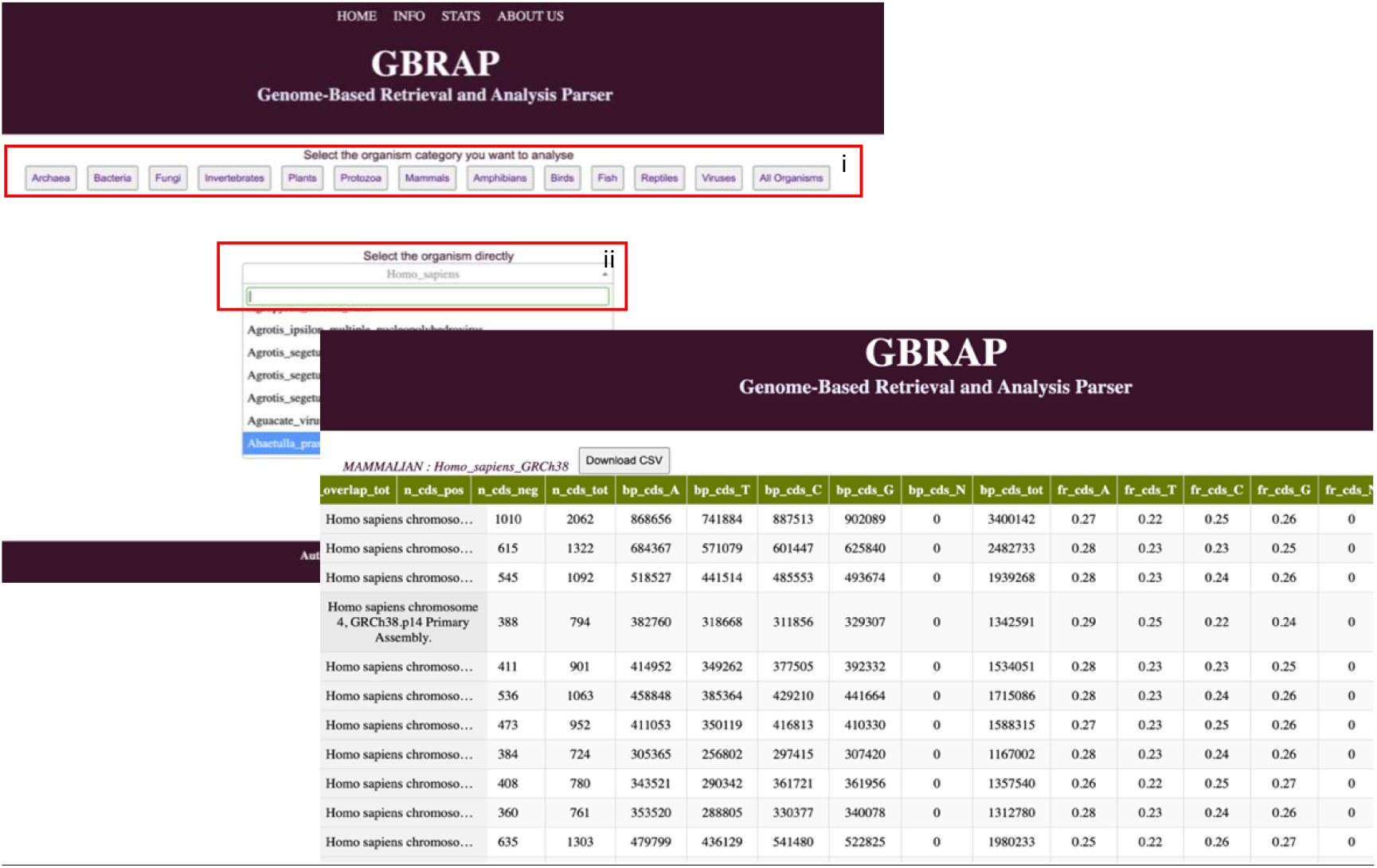
**a)** GBRAP HTML interface with two ways for data extraction: i) Browsing through pre-organized groups or ii) searching directly by the name of the organism. **b)** GBRAP table for Homo sapiens GRCh38 genome showing some statistics of CDS for each chromosome. The first two columns show respectively the count of CDS in the complementary strand and the total in both strands. While next 5 columns show the counts of each of the bases (A, T, C, G) and unknown bases followed by the total length of CDS (bp_cds_tot). The next 5 columns are the base frequencies respectively, followed by the GC content and the remaining columns. (See supplementary material for details about the remaining columns).

Figure 2 depicts a summary of the data included in GBRAP datasets. The first 16 columns of each table contain statistics of the whole chromosome, followed by sets of 20 columns for each genomic element (gene, CDS, intron etc.). The final 64 columns of the table provide codon usage information of the chromosome, which is the frequency of each codon in the coding sequences (see supplementary material for more details about the data columns). All the data are also available for download in comma-separated (CSV) or Excel formats as individual organisms, as combined tables for all organisms in a group, or as one big table containing all the organisms of all groups. Forthcoming updates of the database will empower users to selectively filter data columns or rows as per their requirements.

**Figure 2:**
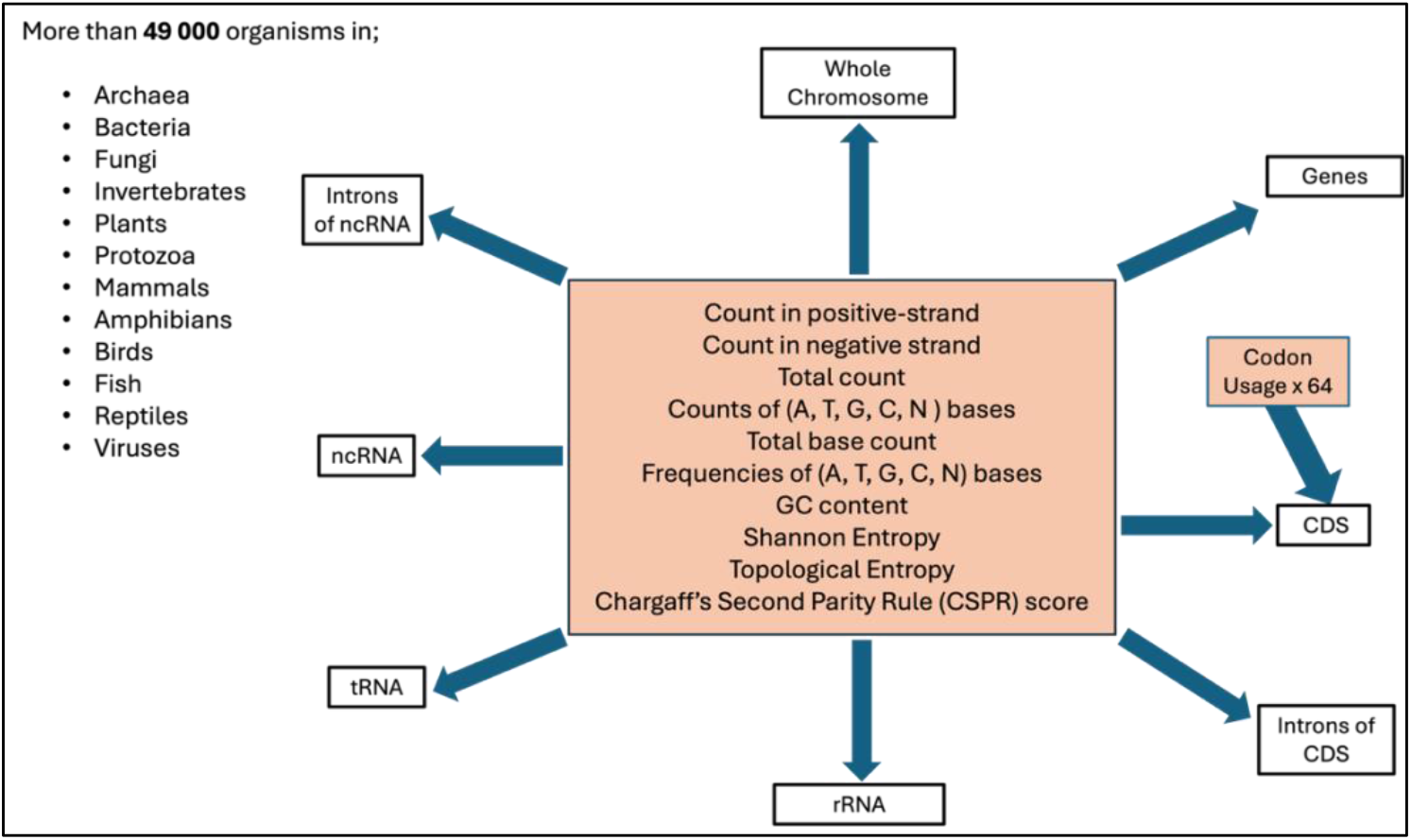
A diagram showing a summary of the data available in GBRAP. All the statistics in the middle box are calculated for the genomic elements shown by arrows. The codon usage of the 64 codons is calculated for coding sequences (CDS). All these data are available for more than 49,000 organisms divided into the listed organism groups.

### Sample Usage

Figure 3 presents some comparisons we performed with GBRAP datasets to highlight the practical usage and reliability of our data. The plots present key genomic features, including genome size, number of genes, GC content, and the most abundant codons across various taxa. With these example graphs, we demonstrate how GBRAP data can be effectively used to draw meaningful conclusions about genomic organization and evolutionary dynamics. For example, the genome size and number of genes provide a broad overview of the genetic complexity across taxa, while the GC content and the distribution of the most abundant codons highlight the evolutionary and functional aspects of the genomes.

**Figure 3:**
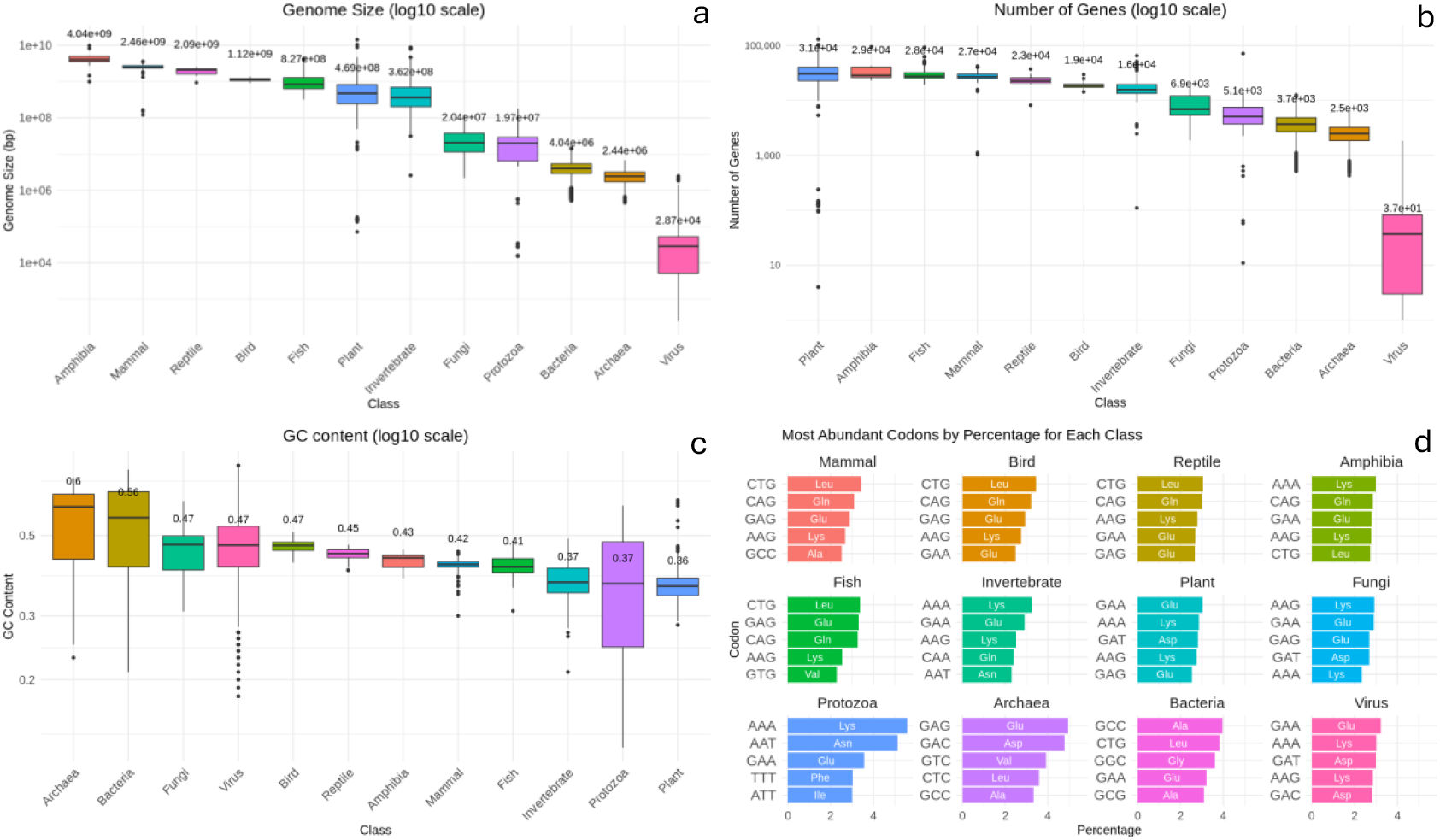
Example graphs to demonstrate the practical usage of GBRAP data. All the graphs show the distribution among different taxonomic groups. (a) Box plot showing genome size distribution (log10 scale). (b) Box plot displaying the distribution of the number of genes (log10 scale). (c) Box plot depicting GC content variation (log10 scale). (d) Bar plot presenting the top 5 most abundant codons.

Additionally, the box plots shown in Figure 3 also highlight the reliability of GBRAP data: panel a) shows the genome size distribution among different taxa which displays that, on average, amphibians and mammals have the largest genome sizes despite some outliers found in the Plant kingdom. Panel b) supports the literature by showing that plants harbour the highest number of genes (Sterck et al. 2007), followed by amphibians, fish and mammals (Wittbrodt et al. 1998). GC content of closely related organisms and of organisms that live in similar environments are found to be similar (Bohlin and H-O Pettersson 2019) which is also seen in panel c) in Birds, Reptiles, Amphibians, and Mammals. Although there are not many articles providing clear information about amino acid abundance in different organisms, some do corroborate the data plotted in panel d), which shows that leucine is a highly expressed amino acid in vertebrates and that closely related species share similarities in codon usage (Nie et al. 2018; Lamolle et al. 2023).

The plots presented in Figure 3 serve as examples of how GBRAP data can be used for descriptive analysis. These observations align closely with established biological principles, thereby validating the accuracy and utility of the GBRAP datasets for research and analysis by the scientific community. Further statistical and analytical work is necessary to confirm the observations presented here and to draw conclusive insights.

### Future Perspectives

As the next step, we are conducting detailed statistical analyses of these data intending to identify new information on evolutionary patterns. Additionally, we are also employing these datasets in various machine-learning analyses. We have recently published one such analysis done using data generated by GBRAP software for Archaea and Bacteria (Bobbo et al. 2024).

We invite the scientific community to utilise the extensive data available through GBRAP database to conduct diverse statistical analyses and machine learning studies to uncover valuable insights and evolutionary patterns among organisms that might not be apparent through conventional analytical methods. We expect that our database will continue to be a valuable resource for the scientific community, enabling further research and innovation across various life domains.

## Acknowledgements

We would like to thank Dr. Marco Ricci for the precious advice on classifying organisms and Dr. Tania Bobbo for her valuable insights on the manuscript. This study has been funded by MAPS Department funds BIRD2021 - prot. BIRD213010, University of Padova, Italy.

## Code and Data Availability

All raw data used in this work were derived from the RefSeq database of NCBI. The Python codes underlying this article are available at <GitHub link will be available here> for the purpose of evaluating the manuscript. It will be made publicly available once the paper has been conditionally accepted.

