## Supplementary material for "GBRAP: A Comprehensive Database and Tool for Exploring Genomic Diversity Across All Domains of Life"

#### GBRAP Overview

A comprehensive software tool to analyse GenBank files and an online database, housing an extensive collection of carefully curated, high-quality genome statistics generated by the software for all the organisms (latest assembly; reference or representative genome) available in the RefSeq database of NCBI. The users can directly search or select from pre-categorized groups, the organisms of their choice and access data tables containing useful genomic information (eg. Base counts, GC content, Shannon Entropy, Chargaff Score) separately calculated for different genomic elements (eg. CDS, Introns, ncRNA) adding up to a collection of more than 200 genomic statistics. The data are separately displayed (if applicable) for each chromosomal, mitochondrial or chloroplast sequence. All the data can be visualised on the database itself or downloaded as CSV or Excel files. The GBRAP database is free to access without any registration and is publicly available at <GBRAP database link will be available here>

#### Installation and Usage

GBRAP is an open-source tool and is freely available at <GitHub link will be available here>. It can be efficiently executed even on a standard laptop without the need for specialized hardware or configurations. Once the user has the gbff file needed to be analysed GBRAP can be used in the command line/ terminal by running the command ". /GBRAP_command_line_tool.py in input_file_name.gbff out output_file_name.csv". GBRAP includes all the needed files for the execution of the complete analysis, and no additional installations are necessary.

GBRAP returns in output a comma-separated file (CSV) that could be imported into excel. Each file contains:

- Locus ID: Sequence ID
- Definition: Sequence description
- bp_chromo_A: Number of 'A' bases in the whole chromosome/genome
- bp_chromo_T: Number of 'T' bases in the whole chromosome/genome
- bp_chromo_C: Number of 'C' bases in the whole chromosome/genome
- bp_chromo_G: Number of 'G' bases in the whole chromosome/genome
- bp_chromo_N: Number of 'N' bases in the whole chromosome/genome
- bp_chromo_total: Total number of bases in the whole chromosome/genome

- fr_chromo_A: Frequency of 'A' bases among total the whole chromosome/genome bases
- fr_chromo_T: Frequency of 'T' bases among total the whole chromosome/genome bases
- fr_chromo_C: Frequency of 'C' bases among total the whole chromosome/genome bases
- fr_chromo_G: Frequency of 'G' bases among total the whole chromosome/genome bases
- fr_chromo_N: Frequency of 'N' bases among total the whole chromosome/genome bases
- GC_chromo: Percentage of 'G' and 'C' bases in the whole chromosome/genome
- chromo_topological_entropy: Topological entropy calculated from the whole chromosome/genome
- chromo_chargaff_pf: Chargaff's second parity rule score (method1) for the whole chromosome/genome
- chromo_chargaff_ct: Chargaff's second parity rule score (method2) for the whole chromosome/genome
- chromo_shannon: Shannon entropy score for the whole chromosome/genome
- n_gene_pos: Number of genes in the positive strand
- n_gene_neg: Number of genes in the negative strand
- n_gene_tot: Total number of genes in the chromosome
- bp_gene_A: Number of 'A' bases in total genes
- bp_gene_T: Number of 'T' bases in total genes
- bp_gene_C: Number of 'C' bases in total genes
- bp_gene_G: Number of 'G' bases in total genes
- bp_gene_N: Number of 'N' bases in total genes
- bp_gene_total: Total number of bases in total genes
- fr_gene_A: Frequency of 'A' bases among total gene bases
- fr_gene_T: Frequency of 'T' bases among total gene bases
- fr_gene_C: Frequency of 'C' bases among total gene bases
- fr_gene_G: Frequency of 'G' bases among total gene bases
- fr_gene_N: Frequency of 'N' bases among total gene bases
- GC_gene: Percentage of 'G' and 'C' bases in total genes
- gene_topological_entropy: Topological entropy calculated from total gene sequences
- gene_chargaff_pf: Chargaff's second parity rule score (method1) for total genes
- gene_chargaff_ct: Chargaff's second parity rule score (method2)for total genes
- gene_shannon: Shannon entropy score for total genes
- bp_gene_overlap_total: Number of bases overlapping between genes on two strands
- n_cds_pos: Number of CDS in the positive strand
- n_cds_neg: Number of CDS in the negative strand
- n_cds_tot: Total number of CDS in the chromosome
- bp_cds_A: Number of 'A' bases in total CDS (coding sequences)
- bp_cds_T: Number of 'T' bases in total CDS
- bp_cds_C: Number of 'C' bases in total CDS
- bp_cds_G: Number of 'G' bases in total CDS
- bp_cds_N: Number of 'N' bases in total CDS
- bp_cds_total: Total bases in total CDS
- fr_cds_A: Frequency of 'A' bases among total CDS bases
- fr_cds_T: Frequency of 'T' bases among total CDS bases
- fr_cds_C: Frequency of 'C' bases among total CDS bases
- fr_cds_G: Frequency of 'G' bases among total CDS bases
- fr_cds_N: Frequency of 'N' bases among total CDS bases
- GC_cds: Percentage of 'G' and 'C' bases in total CDS

- cds_topological_entropy: Topological entropy calculated from total CDS sequences
- cds_chargaff_pf: Chargaff's second parity rule score (method1) for total CDS
- cds_chargaff_ct: Chargaff's second parity rule score (method2) for total CDS
- cds_shannon: Shannon entropy score for total CDS
- bp_cds_overlap_total: Number of bases overlapping between CDS on two strands
- bp_cds_intron_A: Number of 'A' bases in total introns between CDS
- bp_cds_intron_T: Number of 'T' bases in total introns between CDS
- bp_cds_intron_C: Number of 'C' bases in total introns between CDS
- bp_cds_intron_G: Number of 'G' bases in total introns between CDS
- bp_cds_intron_N: Number of 'N' bases in total introns between CDS
- bp_cds_intron_total: Total bases in total introns between CDS
- fr_cds_intronn_A: Frequency of 'A' bases in total introns between CDS
- fr_cds_intron_T: Frequency of 'T' bases in total introns between CDS
- fr_cds_intron_C: Frequency of 'C' bases in total introns between CDS
- fr_cds_intron_G: Frequency of 'G' bases in total introns between CDS
- fr_cds_intron_N: Frequency of 'N' bases in total introns between CDS
- GC_cds_intron: Percentage of 'G' and 'C' bases in total introns between CDS
- cds_intron_topological_entropy: Topological entropy calculated from total intron sequences between CDS
- cds_intron_chargaff_pf: Chargaff's second parity rule score (method1) for total introns between CDS
- cds_intron_chargaff_ct: Chargaff's second parity rule score (method2) for total introns between CDS
- cds_intron_shannon: Shannon entropy score for total introns between CDS
- bp_cds_intron_overlap_total: Number of bases overlapping between introns between CDS on two strands
- n_ncRNA_pos: Number of ncRNA in the positive strand
- n_ncRNA_neg: Number of ncRNA in the negative strand
- n_ncRNA_tot: Total number of ncRNA in the chromosome
- bp_ncRNA_A: Number of 'A' bases in total ncRNA
- bp_ncRNA_T: Number of 'T' bases in total ncRNA
- bp_ncRNA_C: Number of 'C' bases in total ncRNA
- bp_ncRNA_G: Number of 'G' bases in total ncRNA
- bp_ncRNA_N: Number of 'N' bases in total ncRNA
- bp_ncRNA_total: Total bases in total ncRNA
- fr_ncRNA_A: Frequency of 'A' bases among total ncRNA bases
- fr_ncRNA_T: Frequency of 'T' bases among total ncRNA bases
- fr_ncRNA_C: Frequency of 'C' bases among total ncRNA bases
- fr_ncRNA_G: Frequency of 'G' bases among total ncRNA bases
- fr_ncRNA_N: Frequency of 'N' bases among total ncRNA bases
- GC_ncRNA: Percentage of 'G' and 'C' bases in total ncRNA
- ncRNA_topological_entropy: Topological entropy calculated from total ncRNA sequences
- ncRNA_chargaff_pf: Chargaff's second parity rule score (method1) for total ncRNA
- ncRNA_chargaff_ct: Chargaff's second parity rule score (method2) for total ncRNA
- ncRNA_shannon: Shannon entropy score for total ncRNA
- bp_ncRNA_overlap_total: Number of bases overlapping between ncRNA on two strands
- bp_ncintron_A: Number of 'A' bases in total introns between ncRNA
- bp_ncintron_T: Number of 'T' bases in total introns between ncRNA

- bp_ncintron_C: Number of 'C' bases in total introns between ncRNA
- bp_ncintron_G: Number of 'G' bases in total introns between ncRNA
- bp_ncintron_N: Number of 'N' bases in total introns between ncRNA
- bp_ncintron_total: Total bases in total introns between ncRNA
- fr_ncintron_A: Frequency of 'A' bases in total introns between ncRNA
- fr_ncintron_T: Frequency of 'T' bases in total introns between ncRNA
- fr_ncintron_C: Frequency of 'C' bases in total introns between ncRNA
- fr_ncintron_G: Frequency of 'G' bases in total introns between ncRNA
- fr_ncintron_N: Frequency of 'N' bases in total introns between ncRNA
- GC_ncintron: Percentage of 'G' and 'C' bases in total introns between ncRNA
- ncintron_topological_entropy: Topological entropy calculated from total intron sequences between ncRNA
- ncintron_chargaff_pf: Chargaff's second parity rule score (method1) for total introns between ncRNA
- ncintron_chargaff_ct: Chargaff's second parity rule score (method2) for total introns between ncRNA
- ncintron_shannon: Shannon entropy score for total introns between ncRNA
- bp_ncintron_overlap_total: Number of bases overlapping between introns between ncRNA on two strands
- n_tRNA_pos: Number of tRNA in the positive strand
- n_tRNA_neg: Number of tRNA in the negative strand
- n_tRNA_tot: Total number of tRNA in the chromosome
- bp_tRNA_A: Number of 'A' bases in total tRNA
- bp_tRNA_T: Number of 'T' bases in total tRNA
- bp_tRNA_C: Number of 'C' bases in total tRNA
- bp_tRNA_G: Number of 'G' bases in total tRNA
- bp_tRNA_N: Number of 'N' bases in total tRNA
- bp_tRNA_total: Total bases in total tRNA
- fr_tRNA_A: Frequency of 'A' bases among total tRNA bases
- fr_tRNA_T: Frequency of 'T' bases among total tRNA bases
- fr_tRNA_C: Frequency of 'C' bases among total tRNA bases
- fr_tRNA_G: Frequency of 'G' bases among total tRNA bases
- fr_tRNA_N: Frequency of 'N' bases among total tRNA bases
- GC_tRNA: Percentage of 'G' and 'C' bases in total tRNA
- tRNA_topological_entropy: Topological entropy calculated from total tRNA sequences
- tRNA_chargaff_pf: Chargaff's second parity rule score (method1) for total tRNA
- tRNA_chargaff_ct: Chargaff's second parity rule score (method2) for total tRNA
- tRNA_shannon: Shannon entropy score for total tRNA
- bp_tRNA_overlap_total: Number of bases overlapping between tRNA on two strands
- n_rRNA_pos: Number of rRNA in the positive strand
- n_rRNA_neg: Number of rRNA in the negative strand
- n_rRNA_tot: Total number of rRNA in the chromosome
- bp_rRNA_A: Number of 'A' bases in total rRNA
- bp_rRNA_T: Number of 'T' bases in total rRNA
- bp_rRNA_C: Number of 'C' bases in total rRNA
- bp_rRNA_G: Number of 'G' bases in total rRNA
- bp_rRNA_N: Number of 'N' bases in total rRNA
- bp_rRNA_total: Total bases in total rRNA

- fr_rRNA_A: Frequency of 'A' bases among total rRNA bases
- fr_rRNA_T: Frequency of 'T' bases among total rRNA bases
- fr_rRNA_C: Frequency of 'C' bases among total rRNA bases
- fr_rRNA_G: Frequency of 'G' bases among total rRNA bases
- fr_rRNA_N: Frequency of 'N' bases among total rRNA bases
- GC_rRNA: Percentage of 'G' and 'C' bases in total rRNA
- rRNA_topological_entropy: Topological entropy calculated from total rRNA sequences
- rRNA_chargaff_pf: Chargaff's second parity rule score (method1) for total rRNA
- rRNA_chargaff_ct: Chargaff's second parity rule score (method2) total rRNA
- rRNA_shannon: Shannon entropy score for total rRNA
- bp_rRNA_overlap_total: Number of bases overlapping between rRNA on two strands
- ATG, AAG, GTA, etc.: Count of respective codons in CDS.

### Chargaff's second parity rule (CSPR) score meaning

In 1950, Erwin Chargaff discovered that the four nucleotides contained in a DNA double helix (A=Adenine, T=Thymine, C=Cytosine and G=Guanine) are symmetrically abundant in both strands of DNA. This symmetry was called Chargaff's first parity rule. In 1968, Chargaff also discovered that also on each DNA strand, the number of Adenines is almost equal to that of Thymines and the number of Cytosines is almost equal to that of Guanines. The first rule was easily explained by the fact that within DNA strands A matches with T, whereas C matches with G. On the single strand, however, this symmetry (Chargaff's second parity rule) is not easily explained. In 2020 four Italian researchers (Fariselli et al., 2020) discovered that this symmetry is linked to the energy of the DNA molecule which has greater stability when it has a Chargaff's second parity rule score close to one.

##### * Chargaff's second parity rule score calculated using an easy method (PF)

An easy way to calculate Chargaff's second parity rule is this:

$$\mathrm{ABS}\left( {}^{((\#A - \#T)}\!\big/_{(\#A + \#T))} + {}^{((\#C - \#G)}\!\big/_{(\#C + \#G))} \right)$$

where "#" means "number of" and A = Adenines, T = Thymines, C = Cytosines and G = Guanines.

In this way the perfect Chargaff's second parity rule score is zero such as in case of "ATGC". The minimum value depends on the length of the sequence. Python 3 code is as follows:

```
def chargaff_pf(self,sequence):
        '''calculates chargaff score PF'''
        counts = Counter(sequence)

        def safe_pf(x,y):
            if x==0 and y==0:
                return 0
            return abs((x-y)/(x+y))

        a_t=safe_pf(counts['A'], counts['T'])
        c_g=safe_pf(counts['C'], counts['G'])

        PF=a_t+c_g

        return PF
```

##### ** Chargaff's second parity rule score calculated using Cristian Taccioli (CT) method

Chargaff’s second parity rule score calculated using Cristian Taccioli’s method is:

$$\frac{\left(\frac{\#A}{\#T}+\frac{\#C}{\#G}\right)}{2}, where\ \#T\ and\ \#G\ \neq 0$$

$$The\ bases\ with\ the\ highest\ value\ must\ be\ placed\ in\ the\ denominator.$$
$$In\ this\ example\ A\ and\ C\ have\ a\ lower\ value\ than\ T\ and\ G\ respectively.$$

"#" means "number of" and A = Adenines, T = Thymines, C = Cytosines and G = Guanines.

Using this equation Chargaff’s second parity rule score is always between zero and one, where one is the maximum value. For example, the sequence “ATGC” has a perfect score which is one because the number of A is equal to the number of T, whereas the number of C is equal to the number of G. This score does not depend on sequence length. Chargaff’s second parity rule calculated using C.T method can be calculated in Python 3 as follow:

```
def chargaff_ct(self,sequence):
        ''' calculates chargaff score CT '''
        counts = Counter(sequence)

        def safe_ratio(x, y):
            if x == 0 or y == 0:
                return 0
            return min(x, y) / max(x, y)

        a_t_ratio = safe_ratio(counts['A'], counts['T'])
        c_g_ratio = safe_ratio(counts['C'], counts['G'])

        CT = (a_t_ratio + c_g_ratio) / 2
```

### Entropy scores calculation

The concept of entropy was introduced in the early 19th century by Rudolf Julius Emanuel Clausius. It represents a characteristic quantity of the state of a physical system capable of expressing the ability of the system itself to be able to proceed to spontaneous transformations and, consequently, the loss of ability to do work when such transformations occur. In simplified terms, the value of entropy increases when the system undergoes spontaneous variations and therefore loses part of its ability to undergo such variations and perform work. In 1872 Ludwig Boltzmann generalized this concept through the study of statistical mechanics by defining entropy as the degree of disorder of a system. In 1948 Claude Elwood Shannon equated the degree of inaccuracy of a message with disorder. For Shannon, in fact, the entropy of information was the degree of complexity of a message that represents the minimum average number of symbols necessary for the encoding of the message itself.

##### *** Shannon score

The term entropy in information sciences was introduced by Shannon in the paper "A Mathematical Theory of Communication" (Shannon, 1948). Shannon entropy is calculated as follows:

$$H(X) = -\sum_{i=1}^{n} P(x_i) \log P(x_i), \text{ where } P \text{ is the frequency of nucleotides}$$

Python 3 code for Shannon entropy is:

```
def shannon (self,seq,base_counts):
        ''' calculates shanon entropy'''
        nt=base_counts
        if nt[0]==0:
            return ''
        else:
            pA = float(nt[1]/nt[0])
            pC = float(nt[2]/nt[0])
            pG = float(nt[3]/nt[0])
            pT = float(nt[4]/nt[0])

            if pA == 0:
                pA=1
            if pC == 0:
                pC=1
            if pG == 0:
                pG=1
            if pT == 0:
                pT=1

            return                                                       -
((pA*math.log2(pA))+(pT*math.log2(pT))+(pC*math.log2(pC))+(pG*math.log2(pG)))
```

##### † Topological entropy score

Topological entropy is a nonnegative real number that is capable of measuring the complexity of a message. Topological entropy was first introduced in 1965 by Adler, Konheim and McAndrew (Adler et al. 1965). In 2011 Koslicki has defined a new approximation to topological entropy free from the finite sample effects and high dimensionality problems. The formula and code can be retrieved from Koslicki, 2011 (Koslicki et al. 2011).

```
def topology (self,seq):
        '''calculates topological entropy'''
        s= seq.replace("N","") ##Removes the N in the sequence
        n = len(set(s)) # number of bases, this is usually equal to 4
        length = len(s)
        if len(s) != 0 and n > 1:
            logg = math.floor(math.log(length,n))
            neww = (s[:(n**logg + logg)]).lower()
            result_mr = math.log(len(set([neww[i:(i+logg)] for i in range(1,
n**logg+1)])),n)/logg #the topological entropy score
        else:
            result_mr = ''
        return result_mr
```
